# Parent-progeny imputation from pooled samples for cost-efficient genotyping in plant breeding

**DOI:** 10.1101/157883

**Authors:** Frank Technow, Justin Gerke

**Author notes:** These authors contributed equally to this work.

## Abstract

The increased usage of whole-genome selection (WGS) and other molecular evaluation methods in plant breeding relies on the ability to genotype a very large number of untested individuals in each breeding cycle. Many plant breeding programs evaluate large biparental populations of homozygous individuals derived from homozygous parent inbred lines. This structure lends itself to parent-progeny imputation, which transfers the genotype scores of the parents to progeny individuals that are genotyped for a much smaller number of loci. Here we introduce a parent-progeny imputation method that infers individual genotypes from index-free pooled samples of DNA of multiple individuals using a Hidden Markov Model (HMM). We demonstrated the method for pools of simulated maize double haploids (DH) from biparental populations, genotyped using a genotyping by sequencing (GBS) approach for 3,000 loci at 0.125*x* to 4*x* coverage. We observed high concordance between true and imputed marker scores and the HMM produced well-calibrated genotype probabilities that correctly reflected the uncertainty of the imputed scores. Genomic estimated breeding values (GEBV) calculated from the imputed scores closely matched GEBV calculated from the true marker scores. The within-population correlation between these sets of GEBV approached 0.95 at 1*x* and 4*x* coverage when pooling two or four individuals, respectively. Our approach can reduce the genotyping cost per individual by a factor up to the number of pooled individuals in GBS applications without the need for extra sequencing coverage, thereby enabling cost-effective large scale genotyping for applications such as WGS in plant breeding.

## Introduction

With the advent of whole-genome evaluation and other molecular methods in plant breeding [1–3], the ability to generate high volumes of genotype data becomes a critical factor in the success of modern breeding programs [3,4]. Whole genome selection (WGS) [5] in particular is revolutionizing plant breeding programs and strategies [3]. The approach applies whole-genome marker effects parameterized in a fully phenotyped and genotyped estimation population to predict performance from genotype alone in target populations. Accurate selections from genotype facilitate faster and greater genetic gain through shorter cycle lengths and increased selection intensity [6–8]. WGS also opens up new opportunities that were inconceivable previously, such as selection for hybrid performance in the earliest stages of the breeding cycle [9,10] or for performance in yet unobserved environments under strong genotype by environment interaction [11].

WGS creates these possibilities without the need for increased resources for phenotypic testing, but it consequently increases the use of genotype data. Large numbers of genotyped and phenotyped reference individuals are required for building accurate prediction models, in particular to predict performance across generations and unrelated populations in order to shorten breeding cycles [12–15]. To maximize investment return over purely phenotypic selection, WGS should be applied to a large number of genotyped-only target individuals [16]. Application of selection to large numbers of unphenotyped target populations in turn facilitates a massive increase in scale of breeding programs [3], but only in combination with the ability to support the corresponding increase in genotype data. Genotyping costs, even though significantly reduced by technological advances over the last two decades [4,17], therefore remain a critical and limiting factor in implementing a successful WGS strategy [18,19].

Genotype imputation is a promising and well-studied approach to reduce genotyping costs [20]. Imputation of missing genotypes typically relies on linkage disequilibrium generated from shared population history [21,22], genetic linkage due to familial relationships [23,24], or a combination of these forces [25,26]. Many individuals evaluated by modern plant breeding programs are fully homozygous doubled haploid lines (DH) [27,28] derived from biparental crosses between elite inbred parents [29].This system is ideal for parent-progeny imputation, which transfers parental genotype scores to all progeny individuals, each of which may initially carry a much smaller number of genotyped loci. Parent-progeny imputation is recognized as a cost-effective way for generating high resolution marker genotypes for a large number of individuals, particularly in the context of WGS [30–32]

Obtaining genotypes from DNA sequence data, termed genotyping by sequencing (GBS) [17] emerged as another approach to reduce genotyping costs and increase scale. This approach efficiently generates high volumes of genotypic data and holds particular promise for applications in plant breeding and genetics [33–35]. Because GBS methods typically result in a large amount of missing data [36], genotype imputation is an integral component of this technology [33,35,37].

The reduction of costs from the combination of GBS and imputation is limited by the need of a separate sequencing library for each sample. Although many libraries can be multiplexed in a single sequencing run, sample-specific library construction is needed to incorporate a sample-identifying oligonucleotide index. In contrast, methods that do not require individual sample identity achieve cost reduction by pooled genotyping, which combines DNA from several individuals and genotypes them jointly in a single assay [38]. Pooled genotyping provides a cost-effective method to assess allele frequency differences between groups of individuals in order to detect signals of selection [39] or identify loci associated with extreme phenotypes as in bulk segregant analysis [40].Within the context of current GBS approaches, index-free pooling into a single sample eliminates the information needed to link a sequencing read to a unique individual.

Here we develop a method of parent-progeny genotype imputation from index-free DNA samples of two or more individuals to simultaneously reduce both the number of genotyped samples and markers per sample. The method takes advantage of pedigree and linkage information to deduce the genotype probabilities of pooled DH lines relative to their fully genotyped parents. The objective of this study is to provide a proof of concept of this approach using simulated data and to identify variables affecting its accuracy.

## Materials and methods

### Imputation method

Parent-progeny genotype imputation from pooled samples infers the marker locus genotypes of the pooled individuals in reference to the set of their direct ancestors (e.g., the parents of the populations). This requires as input the following four pieces of information

1. the complete marker genotypes of the parents at all loci of interest
2. the genotype of the pooled DNA sample (possibly at only a subset of the marker loci)
3. the pedigree relationship between the pooled individuals and their parents, and
4. the genetic linkage map of the loci

Given this information, we calculate for each locus the posterior probabilities of the identity by decent (IBD) inheritance configurations which describe possible patterns of inheritance from parent to offspring. These probabilities are then used to infer the marker genotypes of the pooled individuals. Hereafter we will use the term ‘imputation’ to indicate the inference of genotype scores of individuals from pooled DNA samples, regardless of whether a marker genotype was observed in the pool or not.

### Introductory example

The following example will introduce the concept intuitively (Figure 1). Assume we are interested in the genotypes of two DH (P_1_ and P_2_) at four biallelic SNP markers (L_1_,…, L_4_). The DH are progeny of two biparental populations with four distinct and fully homozygous inbred lines as parents (I_1_×I_2_ and I_3_× I_4_). The DNA of the two DH is pooled into a single sample and genotyped. The critical task becomes inference, at each locus, of the parent of origin for each DH in the pool. We term the combination of parents of origin the *inheritance pattern* of a locus and denote it as, e.g., I_1_-I_3_.

**Fig 1.**
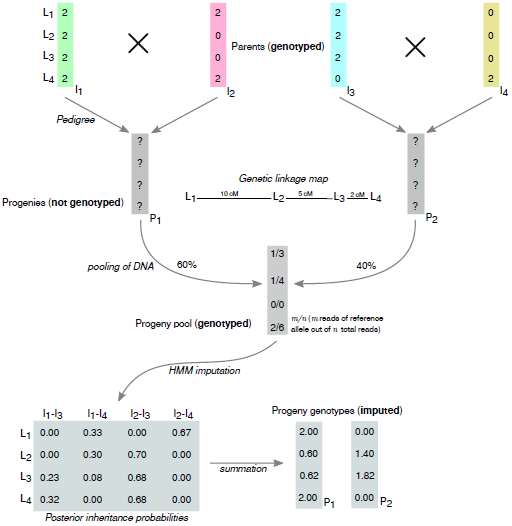
Schematic visualization of parent-progeny imputation. Parent-progeny imputation is carried out for four genetically linked loci L_1_, L_2_, L_3_ and L_4_ for a DNA pool of two DH individuals (P_1_ and P_2_) from two biparental populations (I_1_ × I_2_ and I_3_ × I_4_).

Our example incorporates the read counts of each allele of a marker as would be available if a GBS method is used for genotyping. For simplicity, the parent genotypes at each locus are recoded to represent dosage of a chosen reference allele such that ‘0’ represents a diploid individual with no doses of the reference allele (homozygous alternate), and ‘2’ represents a diploid individual with two doses of the reference allele (homozygous reference). The genotype data for the pool then becomes the sequence read counts of the reference allele relative to the total read count. We will henceforth denote pool genotypes in which only a single allele is present as “homogeneous” and those with multiple alleles as “heterogeneous”, in distinction to homozygous or heterozygous genotypes of individuals. A key factor for inference is the ability to assess whether a pool presents a homogeneous or heterogeneous allelic state at each locus. The technique of inference is therefore not limited to sequencing methods, as any genotyping approach that can detect allelic heterogeneity in the DNA pool would suffice.

In our example, a heterogeneous pool genotype was detected for marker L_1_, with one read of the reference allele out of three total reads. In the absence of genotyping error, the true inheritance pattern must therefore contain both marker alleles. At this locus only parent I_4_ carries the alternate allele and only DH P_2_ can inherit from this parent. Consequently, P_2_ must carry the alternate allele and P_1_ the reference allele. This inference was made possible by knowledge of the parental genotypes and of the pedigree linking parents to DH progeny. A similar reasoning can be applied to locus L_4_ to infer that P_2_ carries the reference allele. A heterogeneous genotype was also detected at locus L_2_. Here, however, the pedigree and genotype information are inconclusive on their own because both the reference and alternate alleles could each be traced to either DH pedigree. For example, the same observed pool genotype could have arisen from P_1_ and P_2_ inheriting respectively from either I_1_ and I_4_ or from I_2_ and I_3_. Although both scenarios are equally likely when locus L_2_ is considered on its own, their relative probabilities can be updated with information from linked loci. Having established the marker genotypes at loci L_1_ and L_4_, and with knowledge of the genetic distance between the loci, it can be shown that the second inheritance pattern (I_2_ and I_3_) is the more likely one because it requires a recombination within a 10cM interval instead of a 7cM interval. Thus, the most likely genotype at L_2_ is the reference allele for P_2_ and the alternate allele for P_1_. We are then left with locus L_3_, for which no read counts were observed for either allele. By combining all of the aforementioned information, it can be shown that again I_2_-I_3_ is the most likely inheritance pattern, because it does not require any additional recombination events beyond the one invoked previously. It follows that P_1_ most likely inherited the alternate allele and P_2_ the reference allele at L_3_. The purpose of this small example was to show how loci with multiple possible inheritance patterns or missing data can be resolved by collectively weighing information from the genetic linkage map, the marker genotypes at linked loci, and the pedigree. Such heuristic reasoning is clearly impractical for more than a few loci and facilitates only very crude inference. A more formal and powerful approach will be described next.

### Parent-progeny imputation with a Hidden Markov Model

If all parents of the pooled DH are present in the ancestor set and the pedigree fully describes all crosses carried out, then the sequence of inheritance patterns along the genomes of the pooled offspring fulfills the requirements of a Hidden Markov Model (HMM). The HMM incorporates the four pieces of information outlined above in the form of the emission and the transition matrix. The emission matrix provides the probabilities that an observed pool genotype could be produced by each possible hidden state of the ancestral inheritance pattern. The transition matrix provides the probabilities that the inheritance pattern at the previous locus can result in a particular pattern at the current locus. These probabilities are a function of both the pedigree and the genetic map. Throughout we assume that the parents of the pooled individuals are fully homozygous inbred lines.

The forward-backward algorithm [41] provides an analytic method to calculate the posterior probabilities of the inheritance patterns for all loci. Given a locus *k*, with an emission matrix ***E***_*k*_, a transition matrix ***T***_*k*_, and a vector of forward probabilities from the previous step (henceforth denoted as ***f***_*k-*1_), the forward pass is

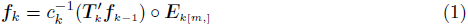

where [*m,*] specifies the row of the emission matrix for the observed genotype *m* (e.g.,*m* counts of a reference allele), ‘o’ refers to element-wise multiplication, and *c*_*k*_ is a normalization constant equal to ((***T**′_k_**f***_*k*-1_) *∘* ***E***_*k*[*m*,]_)′**1**. The backward pass then is

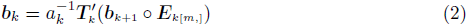

where ***b***_*k*_ indicates the vector of backward probabilities and *a*_*k*_ is similarly defined as *c*_*k*_.

The initial vector of forward probabilities ***f***_0_, which is used when *k* = 1, corresponds to the prior probabilities for the populations involved in the pool. For a pool of two DH from a biparental, F_1_ derived population ***f***_0_ = (0.25 0.25 0.25 0.25)′ (i.e., the products of the expected parental genome contributions to the populations, which are all equal to 0.5 in the case of biparental F_1_ crosses). The initial ***b***_*M*+1_, where *M* is the number of markers, for the backward pass is always ***b***_*M*+1_ = (1 1 1 1)′.

The forward pass is executed from *k* = 1 to *k* = *M* and the backward pass from *k* = *M* to *k* = 1. The posterior inheritance probabilities at locus *k* are then obtained by calculating

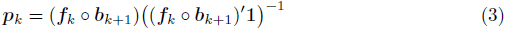

### Transition and emission matrices

We will now use the previously introduced example in Figure 1 to illustrate the derivation of the transition and emission matrices. The transition matrix ***T***_*k*_ for locus *k* for a pool of two F_1_ derived DH from fully homozygous parents is

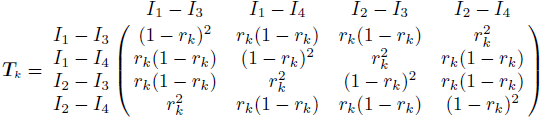

where *r*_*k*_ is the recombination frequency between loci *k* and *k* – 1. For example, the value in row 1 column 2 of this matrix describes the probability that P_1_ inherited from parent I_1_ and P_2_ from parent I_4_ at locus L_2_, conditional on the two progeny inheriting from I_1_ and I_3_, respectively, at locus L_1_. For P_1_ this requires that there is no recombination between the two loci, which happens with probability (1 – *r*_*k*_). For P_2_, the transition from I_3_ to I_4_ requires a recombination event, which has probability *r*_*k*_.Because both events happen independent of each other, the joint probability is *r*_*k*_(1 – *r*_*k*_). The same rationale can be applied to derive transition matrices for different cross types (see S1 File. for BC_1_ derived DH, the second most important cross type in maize breeding, after the F_1_ [29]) or to pooling more than two individuals (see S2 File for a pool of three F_1_ derived DH). Progeny from advanced crosses with additional rounds of meiosis (e.g., F_2_ derived DH or recombinant inbred lines) can also be modeled appropriately. Similar to other parent-progeny approaches, the reduction of linkage between markers in an advanced cross design could lead to lower imputation accuracy unless marker density is increased.

The emission matrix ***E***_*k*_ for locus *k* describes the probability of observing a marker genotype conditional on the inheritance pattern at that locus. The genotype data generated by most sequencing platforms is observed in the form of allele counts and can be modeled with a Beta-Binomial probability distribution. Briefly, the Beta-Binomial distribution models the probability of observing *m* reads of a reference allele out of *n* total reads, when the underlying allele frequency in the sample is uncertain. In principle, this allele frequency is determined by the genotypes of the parents involved in a particular inheritance pattern and can be calculated easily. However, technical variation in quantity and quality of the DNA that each individual contributes to a pool can distort allele frequencies and generate uncertainty [38].

Under the Beta-Binomial model, the probability of observing *m* reference allele reads out of *n* total reads is

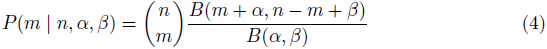

where *B* is the Beta function and *α* and *β* are positive parameters that reflect the uncertainty in the reference allele frequency. The average frequency is given by *α*/(*α* + *β*) and the smaller *α* + *β*, the more variation is expected around it. The parameters were calculated as follows:

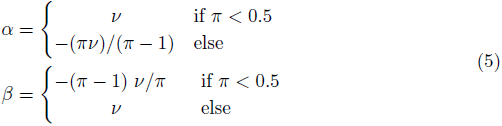

where π is the expected or estimated frequency of the reference allele (with the expected reference allele dosage being *n*π) for a given inheritance pattern and *v* a dispersion parameter reflecting the uncertainty in the estimate. A smaller value for *v* implies greater uncertainty, with *v* > 0 (S1 Fig). We will use *v* = 2 throughout to allow moderate deviation of the allele frequencies from their expected values. A suitable value of *v* in practice can be based upon experimental controls and could be set as locus-specific if desired. The value of π is determined by the genotypes of the parents comprising the inheritance pattern and the proportion of DNA each individual contributed to the pool. This DNA proportion can be estimated from the sequence reads of loci that are monomorphic within each population but for alternate alleles. In the absence of prior estimates, it should be assumed that all individuals contributed an equal amount of DNA. For inheritance pattern I_2_-I_3_ of locus L_2_, for example, π = 0.4, because the reference allele is carried only by I_3_, which would have contributed 40% of the pooled DNA (Figure 1). For inheritance patterns in which all or none of the parents contribute the reference allele, π would be one or zero, respectively. To accommodate small rates of genotyping error or background contamination, the values could be bounded to reflect some uncertainty. For instance, bounds of 0.99 and 0.01 would reflect an expected genotype error rate of 1%. The full emission matrix for L_2_ then would be

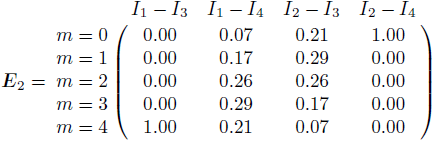

The same principle can be applied to derive emission matrices for different cross types or for pooling more than two DH (see S2 File for the example of a pool of three F_1_ derived DH).

For a genotyping platform that produces categorical genotype calls (i.e., homogeneous reference, homogeneous alternate and the heterogeneous genotype) instead of allele counts, the emission matrix is simply a row vector with a 1 for inheritance pattern that can emit the observed genotype and a 0 for those that cannot. Also in this case, the probability of genotype error could be factored into those values. In case of missing data, such as locus L_3_ in the example, the emission matrix reduces to a row vector of ones, because no data was observed to distinguish among inheritance patterns. In these cases the posterior inheritance probabilities are informed solely by genetically linked loci.

Application of the forward-backward algorithm to the transition and emission matrices for all loci leads to the matrix of posterior inheritance probabilities ***p***_*k*_ shown in Figure 1. A final step is required to convert the posterior probabilities of the inheritance patterns of a locus into imputed marker genotypes. The imputed reference allele dosages of each DH can be calculated by first summing the posterior probabilities of all inheritance patterns containing a parent with the reference genotype and then multiplying by two, i.e.,

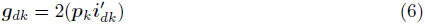

where ***g***_*dk*_ indicates the imputed marker genotype of DH *d* at locus *k* and ***i***_*dk*_ is an incidence vector to indicate the occurrence of the reference allele in the parents of DH *d*. It contains ones to identify inheritance patterns in ***p***_*k*_ for which the relevant parent of DH *d* carries the reference allele and contains zeros where the relevant parent carries the alternate allele.

## Data simulation

We numerically evaluated the described approach using Monte-Carlo simulations of scenarios with varying pool sizes, composition and sequencing coverages. We conducted 1,200 independent replications of each simulated scenario to accurately evaluate the expected values of the statistics of interest, which were then summarized in graphical and tabular form (full results are available in S1 Table, which also includes the standard errors of the estimates). All computations were performed in the R software environment [42].

### Parental inbred line genomes

The simulations were based on the observed genotypes of 35,478 loci with SNP markers of 123 Dent and 86 Flint inbred lines from the maize breeding program of the University of Hohenheim in Germany (the data set is publicly available from the supplement of Technow et al. [8]). The simulated data thus realistically reflects the genome properties such as allele frequency distribution, LD pattern and population structure of this applied maize breeding program, which were described in detail previously [8,43].

### In-silico biparental populations

In each replication of the simulation, we generated in-silico 40 biparental Dent by Dent populations, with random selection of the parents from the set of Dent inbred lines. Each line was restricted to use as a parent of only one population. From each population 25 recombinant DH progeny were generated by simulating meiosis between the loci of the parental lines followed by a chromosome doubling step. This was done with the software package ‘hypred’ [44], which simulates meiosis according to the Haldane mapping function. Together, the Dent populations thus comprised 1,000 recombinant DH. The same procedure was followed to generate 40 Flint populations of size 25.

Simulation of recombination requires a genetic linkage map of the loci. We obtained this by linear scaling of the physical map positions of the loci to the chromosome lengths of the genetic map reported by Fu et al. [45]. This genetic map was subsequently used for parent-progeny imputation, too.

### SNP markers and causal loci

A random sample of 200 loci were considered as ‘causal loci’ of a generic polygenic trait. Those markers were subsequently removed from the set of available loci and treated as unobserved. The causal loci were assigned additive substitution effects drawn from a standard Normal distribution. True genetic values for all DH were then calculated by summing the substitution effects according to the genotypes at the corresponding causal loci. To those we then added a Normal distributed noise variable to generate phenotypic values with heritability of 0.5. The genetic and phenotypic values were used only for later application of WGS. They played no role in the imputation process itself.

As 35,478 loci carry highly redundant information in F1 derived DH families produced by a single generation of meiosis, we randomly selected a subset of 3,000 of the non-causal SNP loci for genotyping and imputation. This number of markers was previously found to be sufficient for WGS in a collection of biparental populations in maize [31]. All subsequent analyses were based on these reduced sets of loci. The true scores of each marker genotype were represented as dosages of the reference allele (i.e., 2 and 0 for the reference and alternate homozygote genotype, respectively). As reference allele in this context we arbitrarily chose the allele with highest allele frequency in the original set of 123 Dent and 86 Flint lines.

### Pooling strategies

We considered pools of two (two-way), three (three-way), and four (four-way) individuals. The pooled individuals either all came from the Dent group (“dent-dent” pools) or from the dent and flint group (“dent-flint” pools). The dent-flint two-way pools comprised one Dent and one Flint individual, three-way pools two Dent and one Flint individual and four-way pools two Dent and two Flint individuals. The pools were formed on a by-population basis, e.g., to form the dent-dent two-way pools, we paired the 25 DH from one Dent population to those of another or to form the four-way pools, we paired the 25 DH from four Dent populations. Within those restrictions, the population pairings and DH pairings within population pairings were chosen at random.

### Simulation of GBS data

To simulate the GBS data of the 3,000 markers for the pooled samples we used the procedure Gorjanc et al. [35] developed for individual samples. The only modification was that we included the possibility of unequal DNA contribution in pooled samples. The step-by-step procedure was as follows

1. Sequenceability of each marker (*seq*_*k*_) was sampled from a Gamma distribution with shape and rate of 4 [35].
2. For a pool *p* the DNA contributions ***d***_*p*_ of the pooled individuals was sampled from a Dirichlet distribution with uniform concentration parameter of 50, 29 or 18 for two-way, three-way and four-way pools, respectively. Those values were chosen such that the standard deviation of each element of ***d***_*p*_ was approximately 0.05.
3. The number of sequence reads *n*_*pk*_ for a pool *p* at marker *k* was drawn from a Poisson distribution with mean *x · seq*_*k*_, with *x* being the targeted sequencing coverage.
4. Finally, the number of reference allele reads *m*_*pk*_ was drawn from a Binomial distribution with success probability equal to sum of the elements of ***d***_*p*_ that correspond to individuals carrying the reference genotype. The number of trials was equal to _*npk*._

As sequencing coverage levels *x* we considered 0.125*x*, 0.25*x*, 0.5*x*, 1*x*, 2*x*, and 4*x*. As in Gorjanc et al. [35], we assumed absence of genotyping errors or DNA contamination. Figure 2 shows how those coverage levels translate into distributions of observed coverages per locus. These values span from the extreme case where data is missing at most marker loci to a more forgiving scenario where coverage is low (typically 1-6 reads) but present for most loci. Even larger values of *x* (and thus higher sequence coverage) would increase accurate detection of heterogeneous pool genotypes. However, since the goal of the approach is to reduce genotyping costs we considered only low coverage scenarios where the resources consumed by parent-progeny imputation from a pooled sample will be competitive with single-sample GBS.

**Fig 2.**
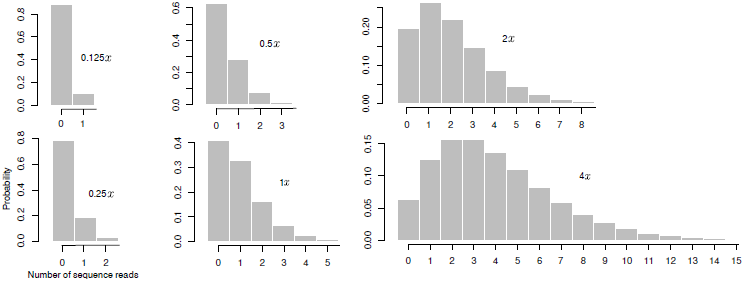
Read number distributions. Read number distribution for different values of targeted sequencing coverage *x*

### GBS cost model

To assess the cost efficiency of pooled genotyping with GBS we used the cost model developed by Gorjanc at al. [35] and available from their supplement. Using the same assumptions for library preparation etc., the cost for genotyping a sample for 3,000 loci at the various sequencing levels were 4*x*: $ 6.20, 2*x*: $ 5.60, 1*x*: $ 5.30, 0.5*x*: $ 5.15, 0.25*x*: $ 5.08 and, 0.125*x*: $ 5.04. To arrive at the genotyping costs per individual, we divided the cost per sample by the number of pooled individuals, assuming that the cost of the pooling step itself was negligible. The ‘true’ marker scores of the DH were treated as obtained from genotyping the individuals separately and at 10*x* coverage. We will henceforth refer to these as high-quality (HQ) marker scores, in contrast to the marker scores obtained from our pooled genotyping approach, which will be referred to as PG marker scores. The cost per individual for the HQ genotyping was $ 8.00.

### Parent-progeny imputation

The HMM was applied to the GBS data to obtain imputed reference allele dosages of the pooled individuals for all 3,000 loci. For this we assumed that the parents were genotyped without error (i.e., it is known without error whether they have the reference or alternate genotype at each locus) and that all genetic positions are known. Locimonomorphic in all populations contributing to a pool provide no information to linked loci and impute with certainty. To reduce computation time we therefore removed monomorphic loci from the HMM and imputed them directly by filling in the scores of the corresponding parents.

### Imputation accuracy

For brevity, imputation accuracy was assessed only for the Dent populations. The pools containing Flint populations were used to measure the effect of pooling more genetically diverse individuals than are observed in a dent-dent pool.

Among the Dent populations, imputation accuracy was measured as the *marker concordance rate* between the true and imputed genotype scores of the polymorphic markers for an individual. We define the marker concordance rate as the percent of markers for which the genotype with highest posterior probability matches the true genotype. Concordance rates were calculated on a by-individual basis and then summarized by the average and standard deviation across individuals. Those statistics were recorded for each replication of the simulation and then averaged across replications, resulting in numerical evaluations of their expected values.

The minimum marker concordance rate depends upon the allele frequency, so concordance should be interpreted relative to a baseline value obtained by a simple imputation of most frequent genotype [46]. In our case this baseline concordance is 50%, because polymorphic markers in biparental populations have an expected minor allele frequency of 0.5.

We further investigated the relationship between the proportion of *multi-polymorphic* markers to total polymorphic markers (*multi-polymorphism rate*) on the marker concordance rate. We defined multi-polymorphic loci as those polymorphic between the parents of at least one more individual in the pool (e.g., locus L_2_ in the example in Figure 1). Because pools were formed on a by-population basis, the proportion of multi-polymorphic markers will be the same for all members of a population. We therefore correlated this rate to the average concordance rate of polymorphic markers in the population. We focused this comparison on the 1*x* coverage level but report results for all other levels in S1 Table.

We also assessed the impact of *imputation uncertainty*, which we define as the posterior probability of the most likely genotype call. As a call becomes more uncertain, the posterior probability will decrease towards the prior for the pool. The average imputation uncertainty was calculated for each individual across all polymorphic loci and across those that were imputed correctly or incorrectly, respectively.

### Assessing impact on WGS

We again evaluated only the Dent populations. A random subset of 30 of the 40 populations was used as the estimation set. As previously mentioned, WGS is most efficient when applied to very large target sets [16]. In our study the target set comprised only the remaining 10 populations, but these can be viewed as representing the performance of a potentially much larger set of target populations. We used the whole genome regression method “BayesB” [5] for estimation of marker effects in the estimation set. This was done with the ‘BGLR’ [47] software package and its default settings for prior distributions and hyperparameters. The BayesB Gibbs-sampler was run for 50,000 iterations. The first 20,000 were discarded as burn-in and only samples from every 3rd subsequent iteration were stored. We used the posterior means as point estimates of the estimated marker effects. These estimates were then applied to the marker scores of the individuals in the target set to produce predictions of their performance in the form of a genomic estimated breeding value (GEBV).

Both the estimation of marker effects as well the calculation of GEBV was done with either the HQ or the PG marker scores. The GEBV obtained when using the HQ scores for estimation and prediction were considered as the “gold-standard” and will henceforth be referred to as “HQ-GEBV”. The GEBV obtained using PG marker scores (for estimation, prediction or both) will collectively be referred to as “PG-GEBV”.

We measured the impact of imputation accuracy and uncertainty on WGS within the target set by calculating the Pearson correlation between the HQ-GEBV and PG-GEBV of the individuals in the target set. We will refer to this measurement as “GEBV concordance”. Thus, whereas the marker concordance rate is a direct measure of imputation accuracy, the GEBV concordance can be understood as measuring it indirectly through its effects on WGS. Other studies investigating the use of imputed marker scores for WGS used the correlation between predicted and true genetic values (commonly referred to as the “prediction accuracy”) as indirect measures of imputation uncertainty [32,35]. We decided against this, however, because the prediction accuracy depends on many other factors that are independent of the genotyping and imputation process, such as the trait heritability or genetic architecture [48]. The GEBV concordance was calculated either across populations (“across GEBV concordance”) or within each population (“within GEBV concordance”). In the latter case the values for the 10 populations were averaged. In each replication of the simulation we further calculated the correlations between the average “within” GEBV concordances and the multi-polymorphism rate of the populations.

## Results and Discussion

Genotype imputation is recognized as an accurate and effective way to reduce genotyping costs for WGS in plant breeding [30–32,34,35,46]. Imputation delivers lower genotype accuracy per sample than could be achieved from fully observed data, but in return it enables larger sample sizes that increase the response to selection and thus the effectiveness of the breeding program overall. In this study we build on this concept of trading small decreases in genotype certainty for large increases in scale by describing a method to genotype two or more individuals from a non-indexed pool of DNA in a single sequencing library. Pooling reduces the per-individual cost of GBS library construction and thereby removes a barrier to genotype cost reduction in low coverage GBS applications [32]. We conducted simulations to investigate the feasibility of pooled sample GBS and varied parameters across different simulated scenarios to assess the impact of sequencing coverage and pool composition on marker and GEBV concordance, and on cost-effectiveness relative to single-sample GBS.

## Marker concordance

We observed generally high concordance rates, with > 95% concordance achieved in two-way pools at 1*x* coverage (Figure 3). Both sequence coverage and pool composition contributed to differences in concordance, with a minimum value near 66% for four-way pools at 0.125*x* coverage, and a maximum near 98% for two-way pools at 4*x* coverage. We will first address the impacts of sequence coverage, then add the variable of pool composition to the discussion.

### The impact of sequence coverage

As expected, increases in sequence coverage improved imputation concordance across all of the coverage rates we tested from 0.125*x* to 4*x* (Figure 3). Concordance increased sharply from the lowest coverage of 0.125*x* until the intermediate coverage value of 1*x* at which point the improvement of concordance leveled off. The strong positive effect of coverage increase on concordance was previously observed for imputation without pooling [35]. Increased coverage can improve imputation through both greater read counts per locus and reduction of the amount of missing data (the number loci represented by zero reads). In our pooled scenario a greater read count at a locus improves the power of inference of the allele dosage, whereas a reduction in missing data increases the information available from linked loci.

The uncertainty stemming from the Binomial sampling process presents a major challenge for allele dosage estimation when *x* is low. At the low sequencing coverage levels used in this study, e.g., *x* < 2, most observed loci are expected to consist of only a single sequence read (Figure 2), which is insufficient to distinguish a heterogeneous from a homogeneous site. Due to the sampling variation inherent in sequencing, the observation of multiple reads still does not guarantee accurate representation of allele dosages. A simple case occurs in a two-way pool with equal sample contribution, where both alleles are expected at equal frequencies. At minimum two reads could accurately capture the allele dosage at such a locus, but under Binomial sampling two reads will still fail to detect heterogeneity 50% of the time. Application of low-coverage GBS to heterozygous or heterogeneous material therefore requires explicit accounting for read sampling uncertainty [35].

We calculate HMM emission probabilities of observed read counts sampled from the underlying allele states according to the Beta-Binomial probability model. This model allows us to account for uncertainty due to sequence read sampling variance. Perhaps more importantly for pooled genotype inference, the π and *v* parameters in the Beta-Binomial model allow an investigator to also account for the uncertainty around the expected allele dosages within the pool (S1 Fig).

Potential sources of uncertainty in allele dosages include unequal quantities of DNA from individual samples and differential amplification of alleles [38]. The parameter π serves to incorporate known or empirically estimated deviations from equal allele dosages. In this study the relative DNA proportions were estimated empirically using read data from loci where the populations generating the pooled individuals were fixed for different alleles (details in methods). For example, a genome-wide π value of 0.6 would represent a 0.6:0.4 ratio of sample DNA contributions in a two-way pool.Differential amplification was absent in our simulations, but it could be measured for each locus from high coverage, non-pooled sequencing of a set of heterozygous individuals [38]. Because differential amplification is locus-specific, incorporation of this source of variation would lead to locus-specific π values. The parameter *v*, which specifies the density of the beta-binomial distribution around π, serves to represent general uncertainty in allele dosages when the deviations cannot be estimated empirically for each locus. For example, *v* could be increased in an experiment expected to generate a greater degree of allele-specific amplification bias. The Beta-Binomial emission model can also incorporate uncertainty due to residual contamination and other sources of genotyping error. To model genotype error for heterogeneous inheritance patterns, *v* can be decreased or increased depending on the amount of genotyping error expected (S1 Fig). Allowance for genotyping error and contamination at homogeneous inheritance patterns must be handled differently. One option is to set maximum and minimum values for the emission probabilities; for example, to a maximum of 0.99 (homogeneous for the expected allele) and a minimum of 0.01 (homogeneous for the unexpected allele) if an error rate of 1% is expected.

Correctly accounting for deviations of allele dosages eliminates their bias but the uncertainty they generate remains, as evidenced by the range of concordance values across pool types and coverage rates. Nonetheless a probabilistic approach enables imputation despite sampling error and low coverage. The concordance rate for two-way pools at 0.125*x* coverage was greater than 80%, suggesting that that many heterogeneous loci are accurately imputed even when represented by a single read. This is possible because the HMM combines sequence read counts at a locus of interest with information from linked loci to jointly calculate the posterior probabilities of each inheritance pattern. This process happens simultaneously for all loci on a chromosome which, in essence, facilitates “borrowing of information” across loci to infer inheritance patterns even with only a small amount of information from each locus. Within this probabilistic framework much of the impact of lower sequence coverage arises from a loss of information from linked loci as more become unobserved. To illustrate this point, we calculated concordance rates for two-way pools in a case where the proportion of missing loci reflected the *x* sequence coverage as before, but the actual observed read counts per locus were capped at a value of 1. In this scenario, greater sequence coverage increases the number of observed loci but provides no additional power to infer the allele dosage at an individual locus. This experiment still displayed a strong increase in marker concordance as the proportion of missing loci decreased, and there was a comparatively small decrease in overall concordance relative to the original simulation that allowed multiple reads per locus (S2 Fig). The result suggests that much of the benefit of increased sequencing coverage comes through the reduction of missing data at linked loci, and this interpretation points to a strategy in which surplus sequencing resources would be better applied to expansion of the number of loci genotyped rather than to increased coverage of a constant set. Our data are derived from simulation, however, and real-world sources of variability such as differential amplification could tip the balance towards increased coverage per locus in order to better inform the parameters of the Beta-Binomial model.

### Number of pooled individuals

We will first discuss the results for the dent-dent pools and later contrast them with the dent-flint pools. The two-way pools resulted in the highest concordance across all coverage rates, followed by three-way and then four-way pools (Figure 3). At the lowest coverage level of 0.125*x*, two way dent-dent pools achieved average concordance around 80%, but three-way pools were instead slightly above 70% and four-way pools slightly below this value (Figure 3). The expected standard deviations of concordance rates from individual to individual for the three and four-way pools were just below 10 percentage points (S3 Fig). This statistic reveals that for a sizable proportion of the individuals the concordance rate was in the vicinity of 50%, which is the baseline value expected from imputation using only population allele frequencies. At 1*x* coverage, the situation improved dramatically. The expected concordance rates for three and four-way pools were at 91% and 87%, respectively, and the standard deviations reduced to 4.4 and 6.5 percentage points. The uncertainty and complexity associated with pooling more than two individuals can thus be largely overcome with a relatively modest increase in coverage.

**Fig 3.**
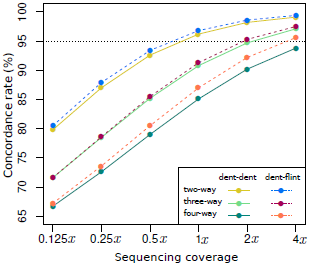
Expected marker concordance rates (%) of polymorphic loci.

An obvious reason for the general decrease in concordance in larger pools is the expansion of possible inheritance patterns. There are only four possible inheritance patterns for a two-way pool, but eight for a three-way and sixteen for a four-way pool. Accurate representation of allele dosages for higher order pools is also more difficult, particularly at low coverage. For example, consider a scenario in which a locus is polymorphic in all populations of a pool. Observation of a single read of the reference allele is sufficient to exclude an inheritance pattern that would emit a homogeneous alternate allele genotype. This would eliminate one of four patterns for a two-way pool, but only one out of eight patterns for a three-way pool and only one out of 16 for a four-way pool. When multiple reads are observed the chance that they capture the true allele dosage is also lower for three and four-way pools because a greater number of more subtle frequency differences must be distinguished. For example, when the true reference allele frequency in a four-way pool is 75%, the chance of actually observing three reference reads out of a total of four is only 42.2%, while the chance of observing four homogeneous reference or alternate reads is still 32%.

The parental allele frequencies also play a role in concordance rates, and their impact can be understood by returning to the hypothetical example in Figure 1. Here locus L_1_ is polymorphic only in the population that generated the second individual P_2_, and this locus is therefore a singly polymorphic locus. In the example, heterogeneous data is observed for this locus, which can only occur with inheritance from parent I_4_. This example shows how singly polymorphic loci can provide strong evidence implicating a specific parent of origin, leading to more certain and accurate imputation. However as the number of individuals in a pool increases, so does the chance that a locus is instead polymorphic in more than one population (multi-polymorphic). As expected, the percent of loci that were multi polymorphic was lowest for two-way pools, followed by three-way and four-way pools (Table 1). On average, higher order pools will contain more multi-polymorphic loci.

**Table 1.** Mean and standard deviation (sd) of the multi-polymorphism rate and its correlation with marker and GEBV concordance

|  | dent-dent |  |  | dent-flint |  |  |
| --- | --- | --- | --- | --- | --- | --- |
|  | two-way | three-way | four-way | two-way | three-way | four-way |
| **multi-polymorphism rate** |  |  |  |  |  |  |
| mean (%) | 39.2 | 61.6 | 74.5 | 21.5 | 52.2 | 60.7 |
| sd (%) | 10.9 | 7.8 | 8.5 | 7.8 | 9.9 | 8.8 |
| **cor. multi-polymorphism rate and marker concordance** |  |  |  |  |  |  |
|  | -0.48 | -0.57 | -0.65 | -0.04 | -0.39 | -0.37 |
| **cor. multi-polymorphism rate and GEBV concordance** |  |  |  |  |  |  |
|  | -0.21 | -0.21 | -0.28 | -0.02 | -0.14 | -0.15 |

### The impact of pool composition

A straightforward objective function for optimizing pool composition would therefore be to choose individuals in a way that minimizes the multi-polymorphism rate. This can be achieved by pooling individuals from populations representing genetically differentiated germplasm groups because allele frequency differences will make it less likely that a locus is polymorphic in multiple populations. The Dent and Flint germplasm groups present two genetically distinct heterotic groups in European maize [8]. Consequently we found that constructing dent-flint pools resulted in a considerably lower multi-polymorphism rate than for the dent-dent pools (Table 1), which translated into a small but consistent increase in marker concordance (Figure 3). The greatest improvements in marker concordance were observed for the four-way pools, which was not surprising given the high multi-polymorphism rate in the dent-dent versions of these pools. The next highest difference, however, was not observed for the three-way pools but the two-way pools. We speculate that this was because moving to the dent-flint version of the three-way pools reduced the Dent germplasm only by one third, whereas it was reduced by half in two and four-way pools. The benefit of the dent-flint arrangement for the three and four-way pools increased with coverage level. In four-way pools, using flint-dent combinations increased the marker concordance rate by more than 1.5 percentage points at 4*x* coverage. The benefit was lower as coverage decreased, but since pooling across germplasm groups does not incur any additional costs relative to pooling within germplasm groups, even small improvements could be worthwhile to pursue. Many of the commercially important field and vegetable crops are bred as hybrid varieties [49,50] that typically target multiple heterotic groups. Pools can be constructed taking advantage of heterotic group divergence in order to optimize singly polymorphic marker rates and thus marker concordance.

Steps can also be taken to promote or avoid pairing of specific populations within a germplasm group. In our study, pools were constructed by randomly selecting populations from within a germplasm group, which led to a high standard deviation for the multi-polymorphism rates among population pairs (Table 1). For example, indent-dent pools the expected standard deviation was 10.9 percentage points around an expected mean of 39.2%. We found that in both dent-flint and dent-dent scenarios the variation in multi-polymorphic rates was negatively correlated with concordance. At 1*x* coverage, the correlation between a population’ s multi-polymorphism rate and marker concordance was strongly negative for most pooling strategies (Table 1). The only exception from this trend were the dent-flint two-way pools, for which the multi-polymorphism rate (21.5%) was very low. Carefully pairing of populations in a way that minimizes the multi-polymorphism rate could therefore result in a further increase in marker concordance. At the very minimum this would involve avoidance of pairing populations that share closely related parents.

### Uncertainty of allele calls

The concordance rate measures the frequency of “erroneous” hard genotype calls. The direct output of the HMM, however, are genotype probabilities which afford a much richer inference that considers the uncertainty around each call. A probability assessment is said to be *calibrated* when an event occurs in p% of the cases in which it was predicted to occur with p% probability. For example, the probabilities from our HMM are calibrated when 80% of the genotype calls made with 80% posterior probability are correct. Figure 4 shows the expected average posterior probability of all genotype calls, for different coverage levels and for the three dent-dent pooling strategies. For example, at 0.125*x* coverage, genotype calls of dent-dent two-way pools were made with 79.2% probability, on average. So we would expect that roughly 79% of them were correct. Comparing this with Figure 3 shows that this was indeed the case, with the corresponding concordance rate being 79.8%. Similarly, at 1*x* coverage the average call probability for two-way pools was 96% and the concordance rate was as well. This close alignment, which holds for all other cases (S4 Fig), shows that the probabilities obtained from the HMM were well calibrated and correctly reflect the uncertainty around each imputed marker score.

**Fig 4.**
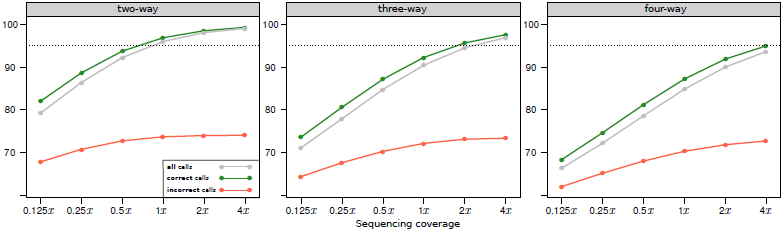
Expected genotype call probabilities (%) for dent-dent pools.

Many applications in statistical genetics, including estimation of whole genome marker effects and calculation of GEBV, do not require hard genotype calls and accept fractional scores proportional to the posterior probability. Carrying over the uncertainty around each marker score into the subsequent analysis, as done in this study, weights each score by the chance of it being incorrect and thus acts as a buffer against imputation error [35]. Indeed, the average certainty of correct calls was always considerably higher than that of incorrect calls (Figure 4). The certainty also increased to a greater degree for correct than incorrect calls, as sequencing coverage increased. Incorrect calls thus not only became fewer but their relative weight in subsequent analyses decreased as well.

## Related computational approaches

Imputation from pooled samples requires assignment of genotype alleles to parental haplotypes, which is a problem also faced when phasing haplotypes in heterozygous individuals. Some phasing algorithms incorporate pedigree information and parent-progeny relationships [26,51], as does pooled genotype imputation. One key difference between the methods is that in a pooled genotyping scenario a genotype can represent more than two haplotypes, as occurs in a three or four-way pool, whereas haplotype phasing is always an attempt to resolve two haplotypes in diploid species.Genome-wide haplotype phasing in polyploid species is considerably more challenging and the methodology is still in its infancy [52]. This potential increase in complexity isalleviated by optimizing the approach specifically for the genetic structure of the populations typically observed in plant breeding. Phasing algorithms are designed to infer haplotypes in populations where all individuals are heterozygous at some loci, such that multiple individuals are often required to accurately infer the phase of the target sample [53]. In plant breeding programs, the parents of populations are in most cases fully homozygous inbred lines genotyped on the full set of marker loci. In this scenario only the parent haplotypes need to be considered to infer the haplotypes of the pooled sample. Further, some phasing approaches, particularly those targeting unrelated individuals, require iterative estimation of haplotype transition probabilities from the data [53]. When a linkage map and pedigree information are available, the transition probabilities can instead be calculated directly. Linkage maps for the 3,000 loci considered here are available [45] and this number of markers was found more than sufficient for genomic selection in biparental breeding populations [31]. Because possible haplotypes are fully represented by the parents and recombination rates do not have to be estimated, imputation is carried out independently for each pool and can easily be parallelized. For example, imputation from the 1,000 dent-flint two-way pools could be carried out on as many CPUs on a high performance computing cluster, with the obvious gains in computing time. Given the very large number of individuals that are generated by modern plant breeding programs [3], this could be an important advantage.

A different method to deconvolute the genotypes of pooled, non-barcoded samples is described by Skelly et al [54] to infer parental origin of homozygous offspring. The distinctions between this approach and ours make each appropriate for different applications. The Skelly et al. approach derives information from the reads that map well to only one of each of a set of parent genomes relative to the other possible parents, which is analogous to using only the singly-polymorphic loci in our approach. The genotypes of each progeny in the pool are deconvoluted individually by modeling a bin-specific read map-ability and Binomial sample of read counts within a bin. An advantage of the read map-ability method is that it does not require a pre-defined set of polymorphic loci. The method does however require sequence information for the parents. The requirement for sequence characterization is an investment justified for populations serving as community resources, but is unrealistic for the breeding scenarios targeted by our approach. A limitation of using uniquely mapping reads is that they cannot inform inheritance in regions of shared ancestry among the parents where only multi-polymorphic loci might be available. Our method jointly models the inheritance of each pooled sample at all loci such that it benefits from alleles unique to a single parent but also leverages information from multi-polymorphic alleles. As we do not rely on read map-ability, our method is better suited to reads containing a low polymorphism rate that does not impact alignment rates. The Skelly et al. method instead takes advantage of reads that will map at different rates across parents, whereas such reads would introduce error into our approach. Our method can therefore be applied to pools of populations with non-sequenced parents that may share ancestry, whereas the Skelly et al. approach is better suited to highly divergent and unrelated parent genomes.

Sonesson et al. [55] demonstrated in a simulation study the use of bulk segregant analysis [40] for estimating whole-genome marker effects from pooled samples. Their approach, however, would require discretizing a continuous trait like grain yield into binary ‘high’ and ‘low’ categories. While this might provide a reasonable approximation for estimating marker effects in some cases, actually using those for WGS would still require the availability of marker genotypes of each selection candidate individually. Their method therefore does not address the main genotyping bottleneck presented by WGS.

### Using array based genotyping platforms

The Binomial sampling error inherent in counting individual sequence reads manifests as the main source of error in allele dosage estimation. Genotyping technologies that do not rely on sequencing, such as fluorescence-based array hybridization, generally achieve less than 5% deviation from the true allele frequency in a sample pool [38]. Other studies indicate that coverages of > 20*x* or perhaps even > 100*x* would be needed to reach this degree of certainty from sequence reads [56,57]. Imputation of pooled offspring provides a less demanding scenario because the possible allele dosages are limited by the number of potential parents in each inheritance pattern (e.g. 0, 0.25, 0.5, 0.75, and 1.0 for a four-way pool). Even with these limited possibilities, distinguishing the correct dosage can be challenging. For example, with a Binomial model of sampling a one-tailed test to distinguish dosages of 2/4 and a 3/4 reference alleles requires 79 reads to achieve 95% power. As discussed previously, in our simulations we achieve high genotype concordance with much lower coverages due to the “borrowing of information” across linked loci. If fluorescence-based array hybridization or other techniques were used for pooled genotyping, then a higher confidence in single allele dosages might lead to comparable imputation concordance with fewer loci overall. As technology currently stands, the need for more marker loci with a sequencing platform is in general outweighed by the lower cost.

## Implications for whole genome selection

In the previous paragraph we discussed the various factors that influence the accuracy of the imputed marker scores and ways to improve it. However, in a WGS scheme, the marker scores themselves are only an intermediate step and matter only in as far as they influence the estimation of marker effects and calculation of GEBV. To assess the impact of the uncertainty added by the imputation, we calculated the “GEBV concordance” as the correlation between PG-GEBV (obtained from imputed marker scores) and HQ-GEBV of individuals in the prediction set.

WGS can be applied within and across populations. Across population selection, however, is largely based on differences in population means [58], which can accurately be predicted from the mean performance of the population parents [59]. The PG-GEBV are expected to reflect differences in population means well, because they are largely the result of differentially fixed alleles, for which imputation in biparental populations is 100% certain. The “across” GEBV concordances were therefore generally considerably higher than their “within” counterparts (S1 Table). Because the real value of WGS in early stages of the breeding cycle comes from the ability to select promising progeny within each population [59], we focused on the “within” GEBV concordance.

Because PG-GEBV are computed from the PG marker scores, factors affecting the marker concordance are expected to have a similar effect on the GEBV concordance. Consequently, the GEBV concordance increased with increasing coverage level and was highest for two-way pools followed by three-way and four way pools (Figure 5). For dent-dent two-way pools, the GEBV concordance was close to 0.60 at the lowest coverage value of 0.125*x* and reached close to 0.95 at 1*x* coverage, when using the PG marker scores for estimation and prediction. For dent-dent four-way pools the corresponding values were considerably lower at 0.30 and 0.69, respectively.

**Fig 5.**
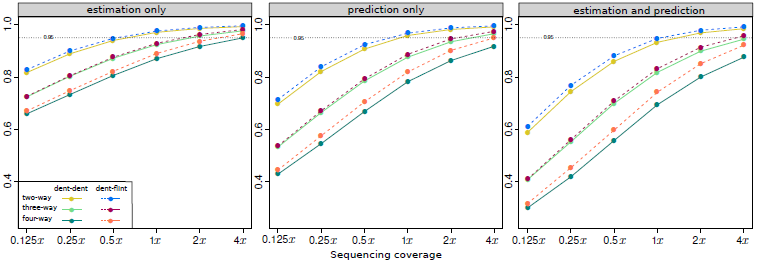
Expected within population GEBV concordance. Imputed marker scores were either used only for estimation of marker effects (“estimation only”), only for calculation of GEBV of target individuals (“prediction only”) or for both (“estimation and prediction”).

Because of the dependence between marker and GEBV concordance, similar optimization options apply. We found that pooling across germplasm groups led to small but consistent increases in GEBV concordance (Figure 5). We also found that the average GEBV concordance of a population was negatively correlated to its multi-polymorphism rate (Table 1), which suggest that pairing individuals in a way that minimizes the multi-polymorphism rate would have a positive effect on the GEBV concordance.

### Scenarios for using imputed marker scores

Three main usage scenarios for the PG marker scores can be distinguished: (1) usage for estimation of marker effects only (“estimation only”), (2) usage for calculation of GEBV in the target set only (“prediction only”) and, (3) usage for both (“estimation and prediction”). The GEBV concordance was higher in the “estimation only” scenario than in the “prediction only” scenario (Figure 5). Estimation of marker effects therefore seems less sensitive than prediction to imputation uncertainty, which was found in other studies as well [32]. Because marker effects are estimated using all individuals in the estimation set, small amounts of imputation error distributed randomly across individuals largely cancel out. If the errors are more concentrated at some loci, for example those with low sequenceability, their effects can be captured by other nearby markers, given the generally high levels of LD observed in plant breeding populations [8,31]. We emphasize again that marker effects were estimated from marker scores proportional to the certainty of the imputed genotype. As we discussed earlier, the weights of erroneously assigned genotypes were considerably closer to the neutral value of 50% (Figure 4), which acted as a buffer against their adverse effects.

GEBVs of individuals in the target set, however, are calculated separately for each individual and after marker effects are estimated. Erroneous marker scores then cannot be compensated for by other individuals or linked markers. Prediction is therefore expected to be more sensitive to the errors and uncertainty introduced by usage of PG marker scores. It is therefore even more important for GEBV calculation than marker effect estimation that the genotype uncertainty be incorporated to lower the impact of loci with a greater chance of being incorrect.

GEBV concordance was lowest when imputed marker scores were used for both estimation and prediction (Figure 5). This was not surprising because of the cumulative effect of the uncertainty and error coming from the estimation and prediction step.

Because WGS is most effective when applied to large numbers of genotyped-only individuals [16], the bulk of the genotyping effort is spent on the target set. The overall cost savings potential of the “estimation only” strategy therefore seems limited in practice. Using PG marker scores for both estimation and prediction has the greatest resource savings potential. However, because the number of individuals in the estimation set is likely going to be small relative to the target set, the difference to the “prediction only” scenario will be small as well and might not justify the increased uncertainty. In addition, the more costly data obtained on the estimation set individuals are their phenotypes. This includes the cost of collecting data in multi-environment field trials for various traits. These costs are still considerable, despite recent advances in high-throughput field phenotyping [60,61]. The genetic values of inbred lines in hybrid crops are evaluated through the performance of their hybrid progeny with multiple testers from the opposite heterotic group [62]. The cost of phenotyping therefore also includes significant costs for producing the testcross seed [63]. It thus seems prudent to maximize the value of the investment in phenotyping by combining it with a high-quality marker genotype, particularly when the individuals in the estimation set are selection candidates themselves [19,64].

This leaves the “prediction only” scenario as the most promising option in practice.Here, the increased genotyping efficiency of parent-progeny impuation from pooled samples is applied to where it matters most: the large numbers of unphenotyped individuals in the target set. For those individuals, the marker genotype is the only investment, apart from the relatively minor cost of creating the inbred line through doubled haploidy or repeated selfing [63]. The vast majority of these individuals will be discarded after their GEBV are assessed. Moderate levels of added uncertainty in the genotypes therefore seem acceptable, if they are overcompensated by increased genotyping efficiency. This trade off will be discussed in the final section.

### Balancing uncertainty and cost efficiency

There is no question that the ability to obtain genotype information of multiple individuals from a single pooled sample considerably decreases genotyping costs. Assuming that the cost of the added DNA pooling step is negligible, genotyping costs would drop two, three, and four-fold, depending on the number of pooled individuals. Additional cost reductions could be achieved by lowering the sequencing coverage level. However, with increasing number of pooled individuals and decreasing sequencing coverage, the GEBV concordance decreases as well, meaning that increased cost savings potential is associated with an increased uncertainty in the calculated GEBV (Figure 5). These two counteracting factors can be balanced by viewing WGS as an indirect selection method and comparing the expected genetic gain when using PG-GEBV or HQ-GEBV as the auxiliary trait.

In general, the standardized response to indirect selection is *R* = *ir*_*A*_*h*, where *i* is the selection intensity on the auxiliary trait, *r*_*A*_ the genetic correlation between the auxiliary and target trait and *h* is the accuracy with which the auxiliary trait can be assessed [65]. In the case of WGS, *r*_*A*_ is the correlation between true and predicted genetic values. For the HQ marker scores *h* = 1, because GEBV can be assessed without error. The indirect selection response for HQ-GEBV thus reduces to *R*_*HQ*_ = *i*_*HQ*_*r*_*A*_. When using PG marker scores, however, the PG-GEBVs themselves are uncertain and so *h*_*P* *G*_ < 1. As estimates of *h*_*PG*_ we used the “prediction only” GEBV concordances (Figure 5) of the dent-dent pools. Because we assumed that PG marker scores were used only for prediction, *r*_*A*_ remains constant. Using pooled genotyping is then expected to be advantageous when the ratio (*i*_*P* *G*_*h*_*P* *G*_)*/i*_*HQ*_ is greater than one.

The selection intensities *i*_*HQ*_ and *i*_*P* *G*_ are calculated from the fraction of selected individuals *s* as *i* = *s*^-1^ϕ(Φ^-1^(1 – *s*)), where ϕ and Φ are the probability density function and cumulative distribution function of the standard Normal distribution, respectively [65]. Let *s*_*HQ*_ and *s*_*PG*_ denote the selected fraction when using HQ or PG marker scores, respectively and let *C*_*HQ*_ and *C*_*PG*_ be the corresponding costs of genotyping a single individual, as obtained from the previously described cost model. Then *s*_*PG*_ = *s*_*HQ*_*C*_*P* *G*_*/C*_*HQ*_, assuming that the same number of individuals is to be selected in each case. For example, if a breeder wants to select 10 individuals from a population and can afford to genotype 50 using the HQ marker platform, *s*_*HQ*_ would be If *C*_*PG*_ is just half of *C*_*HQ*_, 100 individuals could be genotyped with the same resources and *s*_*PG*_ would equal 0.1. As values of *s*_*HQ*_ we choose 0.2 and 0.6. The latter value reflects a scenario in which either WGS is applied only as a pre-test to remove the worst individuals from a population [63] or where the investment per population is low.

The relative merit of PG over HQ increased on average with increasing coverage level (Figure 6). For dent-only two-way pools, it reached a maximum at around 1*x* coverage. It then declined again slowly, as any further increase in *h*_*PG*_, which at this point was already above 0.95 (Figure 5), could not justify the increase in cost. A similar optimum was observed for three-way pools at 2*x* coverage. At very low coverage levels two-way pools had the highest relative merit and four-way pools the lowest, owing to the low GEBV concordance of pools with more than two-individuals. At 1*x* three-way pools had the highest relative merit and finally four-way pools at 4*x* as their GEBV concordance approached 0.95 (Figure 5). The optimal combination of coverage level and number of pooled individuals, i.e., where the relative merit was highest, occurred for the four-way pools at the highest coverage level of 4*x*. Because their GEBV concordance was still notably below 1 at this point, the relative merit did not yet peak, suggesting that the global optimum can be found at even higher coverage levels. This also suggests that the pooling of multiple individuals contributes more to the cost savings potential of the PG approach than the low coverage sequencing per se. The magnitude of the relative merit of PG over HQ at its maximum depended on the initial level *s*_*HQ*_. When *s*_*HQ*_ was low, PG could increase genetic gain by a factor of almost 1.5 over HQ. When it was high (*s*_*HQ*_ = 0.6), the factor was almost 2.5. This is because the selection intensity *i* as a function of *s* follows a curve of diminishing rate of return such that if *s* is low initially, then a much larger decrease in cost and thereby *s* is required to affect a sizable increase in *i*. In such a case it might be advantageous to leave the number of genotyped individuals constant and instead use the freed resources elsewhere. If so, *i*_*PG*_ = *i*_*HQ*_ and the relative merit of using PG marker scores would be equal to the GEBV concordance. We showed that both the marker and GEBV concordance can reach very high values already at intermediate GBS coverage levels, meaning that the penalty in selection gain could be minimal. The tremendous cost savings potential of pooled genotyping could then benefit those components of the breeding operation where the return on investment is greatest.

**Fig 6.**
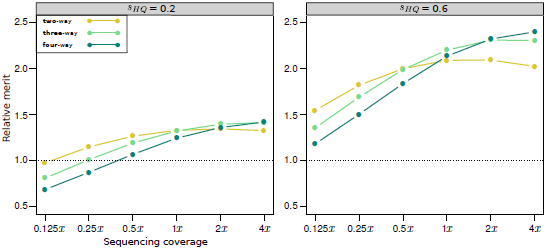
Relative merit of using marker scores imputed from pooled samples for the calculating GEBV in the target set. The merit is expressed relative to expected genetic gain when using the high-quality marker scores for this purpose instead.

To summarize, in this study we presented a method for parent-progeny imputation from pooled samples and applied it to simulated GBS data from biparental populations. We demonstrated that the imputed marker scores can be very accurate even at low coverage levels and then only minimally affect the estimation of marker effects or calculation of GEBV in WGS. The tremendous cost savings potential of the method can therefore facilitate large scale genotyping in plant breeding, a key requirement for successful applications of WGS.

**S1 File. Transition matrix for a pool of two DH derived from a BC_1_ generation** The recurrent and donor parents of the first DH are R_1_ and D_1_, respectively. Those of the second DH are R_2_ and D_2_. The recombination frequency between locus *k* and *k -* 1 is *r*_*k*_.

**S2 File. Example of parent-progeny imputation from a pool of three F_1_ derived DH** Parent-progeny imputation is carried out for four genetically linked loci L_1_, L_2_, L_3_ and L_4_ for a DNA pool of three DH individuals (P_1_, P_2_, P_3_) from three biparental populations (I_1_ × I_2_, I_3_ × I_4_, I_5_ × I_6_).

---

**S1 Fig. Distribution of reference allele dosages under the Beta-Binomial model as a function of *v* and π**

**S2 Fig. Expected marker concordance rate of dent-dent two-way pools when fixing the number of sequence read to one for all observed loci.** The percent of missing markers (in parentheses) correspond to the expectations at the indicated sequencing coverage levels. The full line shows results from the standard GBS scenario where the read number and % missing loci varies as a function of the sequencing coverage *x*. Those results are replicated here for comparison purposes.

**S3 Fig. Expected standard deviation of marker concordance rates.**

**S4 Fig. Average genotype call probability vs. expected marker concordance rate of dent-dent pools**

**S1 Table. Expected marker and GEBV concordances alongside the standard errors of the estimates**

## References

1. Xu Y, Lu Y, Xie C, Gao S, Wan J, Prasanna BM. Whole-genome strategies for marker-assisted plant breeding. Mol Breed. 2012;29: 833–854.

2. MPerez-de Castro A, Vilanova S, Cañizares J, Pascual L, M Blanca J, J Diez M, et al. Application of genomic tools in plant breeding. Curr Genomics. 2012;13: 179–195.

3. Cooper M, Messina CD, Podlich D, Totir LR, Baumgarten A, Hausmann NJ, et al. Predicting the future of plant breeding: complementing empirical evaluation with genetic prediction. Crop Pasture Sci. 2014;64: 311–336.

4. Eathington SR, Crosbie TM, Edwards MD, Reiter RS, Bull JK. Molecular markers in a commercial breeding program. Crop Sci (Supplement 3). 2007;47: S154–S163.

5. Meuwissen THE, Hayes BJ, Goddard ME. Prediction of total genetic value using genome-wide dense marker maps. Genetics. 2001;157: 1819–1829.

6. Heffner EL, Sorrells ME, Jannink JL. Genomic selection for crop improvement. Crop Sci. 2009;49: 1–12.

7. Heffner EL, Lorenz AJ, Jannink JL, Sorrells ME. Plant breeding with genomic selection: gain per unit time and cost. Crop Sci. 2010;50: 1681–1690.

8. Technow F, Schrag TA, Schipprack W, Bauer E, Simianer H, Melchinger AE. Genome properties and prospects of genomic prediction of hybrid performance in a breeding program of maize. Genetics. 2014;197: 1343–1355.

9. Technow F, Riedelsheimer C, Schrag TA, Melchinger AE. Genomic prediction of hybrid performance in maize with models incorporating dominance and population specific marker effects. Theor Appl Genet. 2012;125: 1181–1194.

10. Kadam DC, Potts SM, Bohn MO, Lipka AE, Lorenz AJ. Genomic prediction of single crosses in the early stages of a maize hybrid breeding pipeline. Genes Genom Genet. 2016; p. 3443–3453.

11. Technow F, Messina CD, Totir LR, Cooper M. Integrating crop growth models with whole genome prediction through approximate Bayesian computation. PLoS ONE. 2015;10: e0130855.

12. Jannink JL, Lorenz AJ, Iwata H. Genomic selection in plant breeding: from theory to practice. Brief funct genomics. 2010;9: 166–177.

13. Heffner EL, Jannink JL, Sorrells ME. Genomic selection accuracy using multifamily prediction models in a wheat breeding program. Plant Genome. 2011;4: 65–75.

14. Hickey JM, Dreisigacker S, Crossa J, Hearne S, Babu R, Prasanna BM, et al. Evaluation of genomic selection training population designs and genotyping strategies in plant breeding programs using simulation. Crop Sci. 2014;54: 1476–1488.

15. Schopp P, Müller D, Technow F, Melchinger AE. Accuracy of genomic prediction in synthetic populations depending on the number of parents, relatedness, and ancestral linkage disequilibrium. Genetics. 2017;205: 441–454.

16. Riedelsheimer C, Melchinger AE. Optimizing the allocation of resources for genomic selection in one breeding cycle. Theor Appl Genet. 2013;126: 2835–2848.

17. Elshire RJ, Glaubitz JC, Sun Q, Poland JA, Kawamoto K, Buckler ES, et al. A robust, simple genotyping-by-sequencing (GBS) approach for high diversity species. PloS ONE. 2011;6: e19379.

18. Bernardo R, Yu J. Prospects for genomewide selection for quantitative traits in maize. Crop Sci. 2007;47: 1082–1090.

19. Heslot N, Jannink JL, Sorrells ME. Perspectives for genomic selection applications and research in plants. Crop Sci. 2015;55: 1–12.

20. Li Y, Willer C, Sanna S, Abecasis G. Genotype imputation. Annu Rev Genomics Hum Genet. 2009;10: 387–406.

21. Scheet P, Stephens M. A fast and flexible statistical model for large-scale population genotype data: applications to inferring missing genotypes and haplotypic phase. Am J Hum Genet. 2006;78(4): 629–644.

22. Howie BN, Donnelly P, Marchini J. A flexible and accurate genotype imputation method for the next generation of genome-wide association studies. PLoS Genet. 2009;5: e1000529.

23. Kruglyak L, Lander ES. Faster multipoint linkage analysis using Fourier transforms. J Comput Biol. 1998;5(1): 1–7.

24. Elston RC, Stewart J. A general model for the genetic analysis of pedigree data. Hum Hered. 1971;21(6): 523–542.

25. Meuwissen T, Goddard M. The use of family relationships and linkage disequilibrium to impute phase and missing genotypes in up to whole-genome sequence density genotypic data. Genetics. 2010;185: 1441–1449.

26. O’Connell J, Gurdasani D, Delaneau O, Pirastu N, Ulivi S, Cocca M, et al. A general approach for haplotype phasing across the full spectrum of relatedness. PLoS Genet. 2014;10: e1004234.

27. Longin CFH, Utz HF, Reif JC, Schipprack W, Melchinger AE. Hybrid maize breeding with doubled haploids: I. One-stage versus two-stage selection for testcross performance. Theor Appl Genet. 2006;112: 903–912.

28. Wedzony M, Forster BP, Żur I, Golemiec E, Szechyńska-Hebda M, Dubas E, et al. In: Touraev A, Forster BP, Jain SM, editors. Progress in Doubled Haploid Technology in Higher Plants. Dordrecht: Springer Netherlands; 2009. p. 1–33.

29. Mikel MA, Dudley JW. Evolution of North American dent corn from public to proprietary germplasm. Crop Sci. 2006;46: 1193–1205.

30. Hickey JM, Gorjanc G, Varshney RK, Nettelblad C. Imputation of single nucleotide polymorphism genotypes in biparental, backcross, and topcross populations with a hidden Markov model. Crop Sci. 2015;55: 1934–1946.

31. Jacobson A, Lian L, Zhong S, Bernardo R. Marker imputation before genomewide selection in biparental maize populations. Plant Genome. 2015;8.

32. Gorjanc G, Battagin M, Dumasy JF, Antolin R, Gaynor RC, Hickey JM. Prospects for cost-effective genomic selection via accurate within-family imputation. Crop Sci. 2017;57: 216–228.

33. Poland JA, Rife TW. Genotyping-by-sequencing for plant breeding and genetics. Plant Genome. 2012;5: 92–102.

34. Crossa J, Beyene Y, Kassa S, Pérez P, Hickey JM, Chen C, et al. Genomic prediction in maize breeding populations with genotyping-by-sequencing. Genes Genom Genet. 2013;3: 1903–1926.

35. Gorjanc G, Dumasy JF, Gonen S, Gaynor RC, Antolin R, Hickey JM. Potential of low-coverage genotyping-by-sequencing and imputation for cost-effective genomic selection in biparental segregating populations. Crop Sci. 2017;57: 1–17.

36. Beissinger TM, Hirsch CN, Sekhon RS, Foerster JM, Johnson JM, Muttoni G, et al. Marker density and read depth for genotyping populations using genotyping-by-sequencing. Genetics. 2013;193: 1073–1081.

37. Rutkoski JE, Poland J, Jannink JL, Sorrells ME. Imputation of unordered markers and the impact on genomic selection accuracy. Genes Genom Genet. 2013;3: 427–439.

38. Sham P, Bader JS, Craig I, O’Donovan M, Owen M. DNA pooling: a tool for large-scale association studies. Nat Rev Genet. 2002;3: 862–871.

39. Boitard S, Schlötterer C, Nolte V, Pandey RV, Futschik A. Detecting selective sweeps from pooled next-generation sequencing samples. Mol biol evol. 2012;29: 2177–2186.

40. Michelmore RW, Paran I, Kesseli RV. Identification of markers linked to disease-resistance genes by bulked segregant analysis: a rapid method to detect markers in specific genomic regions by using segregating populations. Proc Natl Acad Sci. 1991;88: 9828–9832.

41. Rabiner LR. A tutorial on Hidden Markov Models and selected applications in speech recognition. Proc IEEE. 1989;77: 257–286.

42. R Core Team. R: A Language and Environment for Statistical Computing; 2014. Available from: https://www.R-project.org/.

43. Technow F, Schrag TA, Schipprack W, Melchinger AE. Identification of key ancestors of modern germplasm in a breeding program of maize. Theor Appl Genet. 2014;127: 2545–2553.

44. Technow F. hypred: Simulation of genomic data in applied genetics; 2013.

45. Fu Y, Wen TJ, Ronin YI, Chen HD, Guo L, Mester DI, et al. Genetic dissection of intermated recombinant inbred lines using a new genetic map of maize. Genetics. 2006;174: 1671–1683.

46. Hickey JM, Crossa J, Babu R, de los Campos G. Factors affecting the accuracy of genotype imputation in populations from several maize breeding programs. Crop Sci. 2012;52: 654–663.

47. de los Campos G, Rodriguez PP. BGLR: Bayesian Generalized Linear Regression; 2016. Available from: https://CRAN.R-project.org/package=BGLR.

48. Zhong S, Dekkers JCM, Fernando RL, Jannink JL. Factors affecting accuracy from genomic selection in populations derived from multiple inbred lines: a barley case study. Genetics. 2009;182: 355–364.

49. Duvick D. Heterosis: feeding people and protecting natural resources. In: Coors J, Pandey S, editors. The genetics and exploitation of heterosis in crops. Madison, WI: CSSA; 1999. p. 19–29.

50. Silva Dias JC. Impact of improved vegetable cultivars in overcoming food insecurity. Euphytica. 2010;176: 125–136.

51. Hickey JM, Kinghorn BP, Tier B, van der Werf JH, Cleveland MA. A phasing and imputation method for pedigreed populations that results in a single-stage genomic evaluation. Genet Sel Evol. 2012;44: 9.

52. Aguiar D, Istrail S. Haplotype assembly in polyploid genomes and identical by descent shared tracts. Bioinformatics. 2013;29: i352–i360.

53. Browning SR, Browning BL. Haplotype phasing: existing methods and new developments. Nat Rev Genet. 2011;12: 703–714.

54. Skelly DA, McCusker JH, Stone EA, Magwene PM. Private haplotype barcoding facilitates inexpensive high-resolution genotyping of multiparent crosses. bioRxiv. 2017;DOI:10.1101/116582.

55. Sonesson AK, Meuwissen TH, Goddard ME. The use of communal rearing of families and DNA pooling in aquaculture genomic selection schemes. Genet Sel Evol. 2010;42: 41.

56. Rellstab C, Zoller S, Tedder A, Gugerli F, Fischer MC. Validation of SNP allele frequencies determined by pooled next-generation sequencing in natural populations of a non-model plant species. PLoS ONE. 2013;8: e80422.

57. Lynch M, Bost D, Wilson S, Maruki T, Harrison S. Population-genetic inference from pooled-sequencing data. Genome Biol Evol. 2014;6: 1210–1218.

58. Windhausen VS, Atlin GN, Hickey JM, Crossa J, Jannink JL, Sorrells ME, et al. Effectiveness of genomic prediction of maize hybrid performance in different breeding populations and environments. G3. 2012;2: 1427–1436.

59. Riedelsheimer C, Endelman JB, Stange M, Sorrells ME, Jannink JL, Melchinger AE. Genomic predictability of interconnected biparental maize populations. Genetics. 2013;194: 493–503.

60. Montes JM, Technow F, Dhillon BS, Mauch F, Melchinger AE. High-throughput non-destructive biomass determination during early plant development in maize under field conditions. Field Crop Res. 2011;121: 268–273.

61. Araus JL, Cairns JE. Field high-throughput phenotyping: the new crop breeding frontier. Trends Plant Sci. 2014;19: 52–61.

62. Albrecht T, Wimmer V, Auinger HJ, Erbe M, Knaak C, Ouzunova M, et al. Genome-based prediction of testcross values in maize. Theor Appl Genet. 2011;123: 339–350.

63. Longin CFH, Mi X, Würschum T. Genomic selection in wheat: optimum allocation of test resources and comparison of breeding strategies for line and hybrid breeding. Theor Appl Genet. 2015;128: 1297–1306.

64. Müller D, Technow F, Melchinger AE. Shrinkage estimation of the genomic relationship matrix can improve genomic estimated breeding values in the training set. Theor Appl Genet. 2015;128: 693–703.

65. Falconer DS, Mackay TFC. Introduction to quantitative genetics. 4th ed. Harlow, UK: Pearson; 1996.

